# Data Distribution for Phylogenetic Inference with Site Repeats via Judicious Hypergraph Partitioning

**DOI:** 10.1101/579318

**Authors:** Ivo Baar, Lukas Hübner, Peter Oettig, Adrian Zapletal, Sebastian Schlag, Alexandros Stamatakis, Benoit Morel

## Abstract

The so-called site repeats (SR) technique can be used to accelerate the widely-used phylogenetic likelihood function (PLF) by identifying identical patterns among multiple sequence alignment (MSA) sites, thereby omitting redundant calculations and saving memory. However, this complicates the optimal data distribution of MSA sites in parallel likelihood calculations, as the cost of computing the likelihood for individual sites strongly depends on the sites-to-cores assignment. We show that finding a ‘good’ sites-to-cores assignment can be modeled as a hypergraph partitioning problem, more specifically, a specific instance of the so-called judicious hypergraph partitioning problem. We initially develop, parallelize, and make available HyperPhylo, an efficient open-source implementation for this flavor of judicious partitioning where all vertices have the same degree. Using empirical MSA data, we then show that sites-to-core assignments computed via HyperPhylo are substantially better than those obtained via a previous na ï ve approach for phylogenetic data distribution under SRs.

## I. INTRODUCTION

Phylogenetic inference, that is, the reconstruction of evolutionary trees based on the molecular sequence data of the species under study, has numerous applications in medical and biological research. Thus, tools for phylogenetic inference such as RAxML [1] or MrBayes [2] are widely used and highly cited. With the advent of new molecular sequencing technologies, the field faces a substantial scalability challenge as phylogenetic analyses of empirical data sets under the standard likelihood models of sequence evolution can require thousands to millions of CPU hours. Most state-of-the art tools for phylogenetic inference have already been parallelized and are regularly deployed on large clusters for production runs. Therefore, an optimal data distribution is key for achieving ‘good’ parallel efficiency.

Here, we analyze specific properties of the, so far, most complex variant of the phylogenetic data distribution problem. We initially transform the problem into a hypergraph partitioning problem and show that it corresponds to a particular instance of the so-called judicious hypergraph partitioning problem [3] (henceforth, denoted as judicious partitioning). As no efficient implementation of a judicious partitioning algorithm was available, we develop, parallelize, and present HyperPhylo^1^ an efficient open-source implementation of the specific judicious partitioning flavor where all vertices have the same degree. We then apply HyperPhylo to our phylogenetic data distribution problem and compare the data distribution computed via judicious partitioning to the data distribution obtained via a simple ad hoc heuristic [4]. We find that phylogenetic data distributions computed with HyperPhylo are substantially better than those obtained via the ad hoc heuristic.

The remainder of this paper is organized as follows. In Section II we introduce the phylogenetic preliminaries, motivate as well as state our problem, and discuss related work. In the subsequent Section III we introduce judicious partitioning and describe our efficient algorithm, implementation, and parallelization. In Section IV we present experimental results. We conclude in Section V.

### II. PHYLOGENETIC PRELIMINARIES

The input to a phylogenetic inference is a *multiple sequence alignment* (MSA) which comprises the sequences of the species under study. The MSA columns are also called *sites*. In empirical phylogenetic analyses, the sites of the MSA are typically subdivided into *p* disjoint *partitions* (e.g., corresponding to individual genes) for which a separate set of likelihood model parameters is estimated. Given the MSA and a partitioning scheme, one can calculate the phylogenetic likelihood function (PLF) on a given candidate tree. PLF calculations typically account for 85-95% of total running time in current tools such as RAxML and MrBayes. Thus, the PLF is *the* target function for optimization and parallelization. PLF calculations are typically parallelized over MSA sites as persite likelihood calculations are independent from each other. Once all per-site likelihoods have been computed, only one collective communication is required to compute the product (or the sum over the logarithms) over all per-site likelihoods to obtain the overall likelihood score. For instance, if there is only one partition, the MSA sites can be distributed to the cores in a cyclic fashion. However, given a list of *p* partitions and *c* cores for a partitioned MSA, finding a ‘good’ (w.r.t. load balance) site-to-core assignment becomes more challenging [5]. As the per-site likelihood calculation cost is identical for all sites, each partition has a computation cost that is linear in the number of sites it comprises.

At an abstract level, the data distribution problem can be stated as follows: For a given number of *c* cores we need to balance the data (MSA sites) among the *c* cores such that the maximal per-core load (sites assigned to core) is minimized. The MSA partitions *are* divisible, since a partition consists of sites whose likelihoods can be computed independently. We can thus improve load balance by splitting partitions into disjoint sets (hereafter referred to as *blocks*) of sites that are allocated to distinct cores. However, splitting a partition does not come for free as the computational cost of a partition has two components: All partitions incur an identical constant base cost *α* at every core (see [5] for details). Thus, if we split a partition among cores, each block of that partition incurs this base cost *α*. For instance, if the sites of a partition are assigned to two distinct cores, *α* needs to be computed redundantly (i.e., once per core). The second component of the per-partition PLF calculation cost is the variable cost *β* which is linear in the number of sites per partition. To maximize parallel efficiency, we need to minimize redundant calculations of *α* by only splitting partitions when necessary, while distributing sites evenly among cores. In [5], we have shown that the problem is NP-hard and introduced an approximation algorithm which solves it in practice.

Thus far, we assumed that per-site calculations costs are identical for each site. That means, if we split a partition into two blocks of equal size among two cores, the respective computational cost is *α* + *β/*2 per core. With the introduction of *site repeats* (SR [6]), an algorithmic optimization of the PLF, the above does not hold. The technique takes repeating (shared) patterns in distinct partial sites (subsets of characters of MSA sites) of a partition into account to reduce persite calculation cost. The amount of savings depends on the current tree topology *and* the MSA. Site repeats reduce the cost *β* by detecting and re-using identical intermediate results among two or more sites belonging to the same partition. Thus, distinct sites of a single partition now exhibit *varying* computational cost. If we assign two sites that share a large fraction of common intermediate results to different cores, the accumulated computational cost and memory for those two sites will increase.

The above complicates data distribution of parallel SR-based PLF calculations. We can still split a single partition among several cores, but now have to sacrifice some SRs which induces additional redundant computations. We denote this phenomenon as *repeat loss*. Note that, computing a lower bound for the minimal repeat loss is straight-forward, as the minimum amount of SR-based calculations can easily be computed for a sequential execution of the PLF (see [7]). Thus, for SR-based parallel PLF computations we might have to also conduct redundant computations for component *β*. Hence, the cost *β* increases if sites that belong to the same partition and share repeats are allocated to distinct cores [7]. We illustrate the impact of SRs on load balance via a simple example (Figure 1). Given a tree, assume a single MSA partition comprising 5 sites that has to be split among 2 cores, and contains some SRs (highlighted by colored boxes). In Figure 1 we depict the required PLF calculations in terms of so-called Conditional Likelihood Vector (CLV) updates—involving a substantial amount of floating point operations—per core with and without SRs, respectively.

**Fig. 1.**
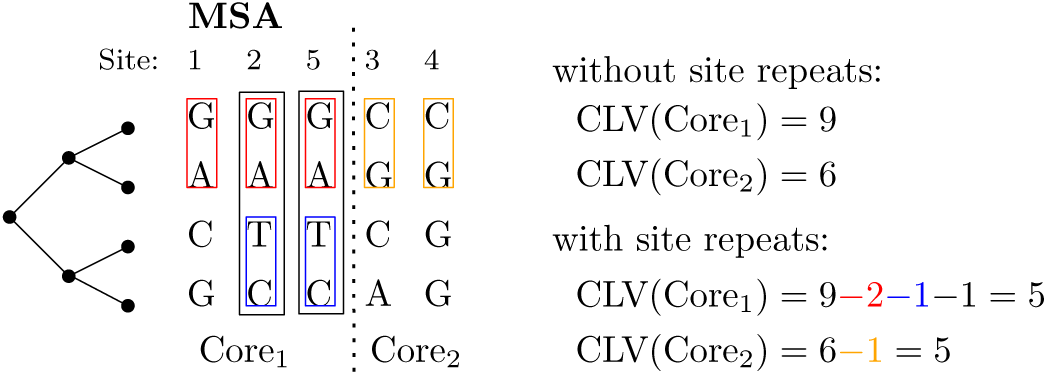
One MSA partition split among two cores and its respective PLF computation costs with and without SRs.

Our goal is thus to devise a data distribution algorithm that yields a ‘good’ trade-off between minimizing repeat loss and load balance in terms of floating point operations per core to, in turn, reduce overall parallel PLF execution times.

We initially outline our current ad hoc randomized data distribution algorithm (RDDA [4]) and then formally state the problem. Given a MSA, and a partitioning scheme, RDDA initially computes the fraction of expected SR-induced computational savings for each partition over a set of random trees. We have empirically assessed that using random trees is sufficient to account for the variance of topology-dependent SR-induced savings. RDDA then simply distributes the partitions according to these fractions using the algorithm from [5]. As RDDA is unaware of repeat loss, that is, sites that share a large proportion of repeats *can* be assigned to different cores. Hence, the obtained data distribution can become sub-optimal. To the best of our knowledge, the RDDA algorithm and our previous exploratory work [7] constitutes the only related work on this problem.

For the remainder of this paper, we will assume that (i) we are given a fixed phylogenetic tree and (ii) that we only have *one* MSA partition (i.e., *p* := 1) whose sites we need divide into two or more blocks in order to assign them to two or more cores, since the coarse-grain distribution (i.e., assigning *entire* MSA partitions to cores) is already handled well by RDDA. To now formally state our problem, we introduce some definitions.

Let *S* be a set of *n* MSA sites and *c* the given number of cores. By 𝒜_*c*_(*S*) we denote the set of all possible sites-to-cores assignments to *c* cores (i.e., all possible partitions of *S* into *c* blocks). An element *Z ∈* 𝒜_*c*_(*S*) then represents a specific sites-to-cores assignment where |*Z*| = *c*.

Let *flops*(*ζ*) be the number of accumulated floating point operations required to compute the per-site likelihoods of all sites *ζ* assigned to a core for one specific sites-to-cores assignment *Z* in 𝒜_*c*_(*S*). Note that it is straight-forward to exactly compute *flops*(*ζ*) by calculating the SRs of the sites in *ζ*, without having to carry out the actual compute-intensive likelihood calculations.

Given these definitions, we can now state the simple problem of splitting just *one* (*p* := 1) MSA partition among *c* cores. Find the sites-to-cores assignment *Z* ∈ 𝒜_*c*_(*S*) that minimizes:

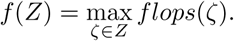

## III. JUDICIOUS HYPERGRAPH PARTITIONING

### A. Preliminaries

An *unweighted, undirected hypergraph H* = (*V, E*) is defined as a set of *n* vertices *V* and a set of *m* hyperedges/nets *E*, where each net *e* is a subset of the vertex set *V* (i.e., *e* ⊆ *V*). The vertices of a net are called *pins*. A vertex *ν* is *incident* to a net *e* if *ν* ∈ *e*. I(*ν*) denotes the set of all incident nets of *ν*. The *degree* of a vertex *ν* is *d*(*ν*) := |I(*ν*)|. With Δ, we denote the maximum degree of a hypergraph. The *size* |*e*| of a net *e* is the number of its pins. A *k-way partition* of a hypergraph *H* is a partition of its vertex set into *k blocks* Π = {*V*_1_, *…, V*_*k*_} such that 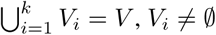 for 1 *≤ i ≤ k*,and *V*_*i*_ *∩ V*_*j*_ *= Ø* for *i ≠ j.* Given a *k*-way partition Π, the number of pins of a net *e* in block *V*_*i*_ is defined as Φ(*e, V*_*i*_) := |{*ν ∈ V*_*i*_ |*ν ∈ e*}|. Net *e is connected* to block *V*_*i*_ if Φ(*e, V*_*i*_) *>* 0.

*Judicious hypergraph partitioning* (JDP) strives to find a *k*-way partition Π of a hypergraph *H* that minimizes the *maximum* number of nets a block is connected to. In other words, judicious partitioning attempts to minimize max(*L*(*V*_1_), *…, L*(*V*_*k*_)), where *L*(*V*_*i*_) := |{*e ∈ E* | Φ(*e, V*_*i*_) *>* 0}| is the load of a block *V*_*i*_. The problem is known to be NP-hard [8] and has mainly been studied in the context of extremal combinatorics [9]–[11]. To the best of our knowledge, Tan *et al.* [3] were the first to investigate algorithmic aspects of the problem.

JDP differs significantly from the classical *k*-way hypergraph partitioning problem (HGP). While JDP is *solely* concerned with minimizing the *maximum* number of nets a block is connected to, the goal of HGP is to partition the vertex set into *k* disjoint blocks of *bounded size* (at most 1 + *ε* times the average block size) while minimizing an objective function such as cut-net (i.e., the weight of all nets that connect more than two blocks) or connectivity (which additionally takes into account the actual number of blocks connected by a cut net). Because of this difference, we do not use existing HGP tools such as hMetis [12] and KaHyPar [13], which are geared towards computing vertex-balanced partitions with small cuts.

### B. Connection between Judicious Partitioning and the Phylogenetic Data Distribution Problem

The input of the phylogenetic data distribution problem is one MSA partition with *n* sites *s*_*i*_, a binary tree topology specifying the order of PLF calculations (and hence, the SRs), and a given number of cores *c*. Two MSA sites *s*_*i*_ and *s*_*j*_ are repeats of one another at an inner node *q* of the phylogenetic tree, if the partial sites *s*_*i*_ | *q* and *s*_*j*_ | *q* induced by the subtree rooted at *q* are *exactly* identical (e.g., the colored boxes in Figure 1 are such partial sites). By partial site, we denote those nucleotides of a MSA site that are contained in a subtree rooted at *q*. If two partial MSA sites *s*_*i*_ | *q* and *s*_*j*_ | *q* are repeats of one another we say that they belong to to the same *repeats class*. Thus, each partial site will be in exactly one repeats class at every inner node of the phylogenetic tree. Note that if a site does not repeat any other site at an inner node, it is assigned to a separate repeats class of size 1. At the root of our example phylogeny in Figure 1, core 1 has two repeat classes (sites *{*1*}, {*2, 5*}*) and core 2 also has two repeat classes (sites *{*3*}, {*4*}*). The number *flops*(*ζ*) of accumulated floating point operations required to compute the per-site likelihoods of all sites *ζ* assigned to a core is directly proportional to the accumulated number of repeats classes over all inner nodes of the given phylogenetic tree. For instance, the number of accumulated repeats classes for core 1 in our example is 1 + 2 + 2.

We can now use a hypergraph *H* = (*V, E*) to formulate our data distribution problem: Each site *s*_*i*_ corresponds to a vertex *ν*_*i*_ *∈ V* and each repeats class *r* = *{s., …, s.}* induced by an inner node of the phylogenetic tree corresponds to a hyperedge *e ∈ E* of size |*r*| containing the corresponding vertices *e* = {*ν., …, ν.*}. Thus, the number of vertices in the hypergraph is equal to the number of sites of the MSA and the number of hyperedges corresponds to the total number of repeats classes induced by all inner nodes of the entire tree topology. An example is shown in Figure 2. Since each inner node of a phylogenetic tree incurs computations for all MSA sites, each site is in *exactly one* repeat class for each inner node of the phylogeny. Therefore, each inner node incurs one hyperedge for each MSA site and the degree of each vertex *ν*_*i*_ equals the number of repeats classes the corresponding site *s*_*i*_ belongs to. By construction, the degree of every vertex *ν ∈ V* is equal to the number of inner nodes of the phylogeny, that is, *d*(*ν*) = Δ, *∀ν ∈ V*.

**Fig. 2.**
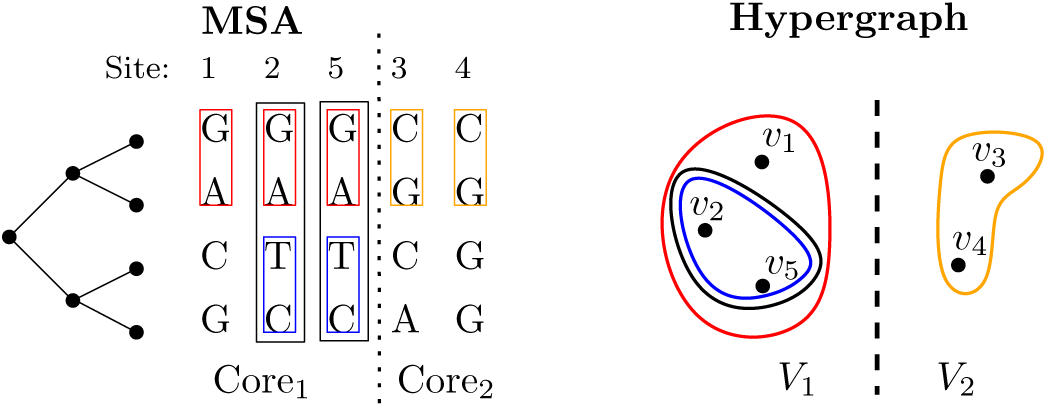
Example of a partitioned MSA and the corresponding partition Π = *{V*_1_, *V*_2_*}* of the hypergraph *H*. Nets of size 1 have been omitted for clarity.

The site-to-cores assignment *Z* ∈ 𝒜_*c*_ that minimizes max_*ζ∈Z*_ *flops*(*ζ*) for a given number of cores *c* therefore corresponds to a judicious *k*-way partition of the corresponding hypergraph *H* into *k* = *c* blocks.

*A note on terminology:* In phylogenetic inference, a partition of a MSA corresponds to a subset of sites (e.g., corresponding to individual genes), whereas in graph and hypergraph partitioning literature the term partition is used in the mathematical sense, that is, a partition is a grouping of the vertex set into non-empty, pairwise-disjoint subsets called blocks. In the following, we therefore explicitly use the term *MSA partition* to refer to a subset of sites of a MSA.

### C. The Judicious Partitioning Algorithm of Tan et al. [3]

The algorithm is based on the insight that the objective score, that is, the cost of a *k*-way partition Π of a hypergraph *H* can be bounded both, from below, and from above. The lower bound is obtained by observing that the maximum load *L*(*V*_*i*_) of a block *V*_*i*_ cannot be smaller than the maximum degree Δ. This is because the maximum-degree vertex has to be assigned to one of the blocks. An upper bound for *L*(*V*_*i*_) is given by the number of hyperedges |*E*|.

The algorithm does not directly partition the hypergraph into *k* blocks while minimizing the maximum load. Instead, it repeatedly computes partitions of increasing cost Δ + *d* for0 *≤ d ≤* |*E*| *-* Δ without restricting the number of blocks a priori. Once the number of blocks *k*′ of a computed partition is ≤ *k*, the algorithm stops and returns the current partition. Otherwise (i.e., if *k*′ ⪈ *k*), the algorithm uses the current solution as input for the next round/iteration. Thus, it increases the cost in each round to find a partition into fewer blocks.

To compute a partition with a solution cost of Δ + *d* the algorithm relies on solving a minimum set cover problem. In the first pass of the algorithm (i.e., *d* = 0) the set cover problem is constructed as follows: The universe 𝒰 is defined to be the set of vertices. Each vertex *ν* is implicitly represented via its incident nets I(*ν*), that is, vertex *ν*_*i*_ is encoded as I(*ν*_*i*_). Let 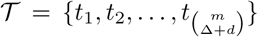 be the set that contains all combinations for choosing Δ + *d* out of *m* hyperedges. The collection 𝒮 of sets that cover *U* is then defined as 𝒮 = 𝒮 {𝒮_1_, 𝒮_2_, *…*}, where 𝒮_*j*_ = {I(*ν*) | (*ν*) *⊆ t*_*j*_, *ν* ∈ *V*}. Then, the standard greedy algorithm (i.e., in each round, choose the set that contains the largest number of uncovered elements) is used to find a sub-collection 𝒮 *\** of 𝒮 that covers 𝒰. In later rounds of the algorithm, 𝒰 is replaced by the current solution 𝒮 *\**. In each iteration, 𝒮 *\** thus induces a partition of the elements in 𝒰, which in turn corresponds to a partition of the vertex set of the hypergraph. We provide the corresponding pseudo code in Algorithm 1. An example is shown in Figure 3.

**Fig. 3.**
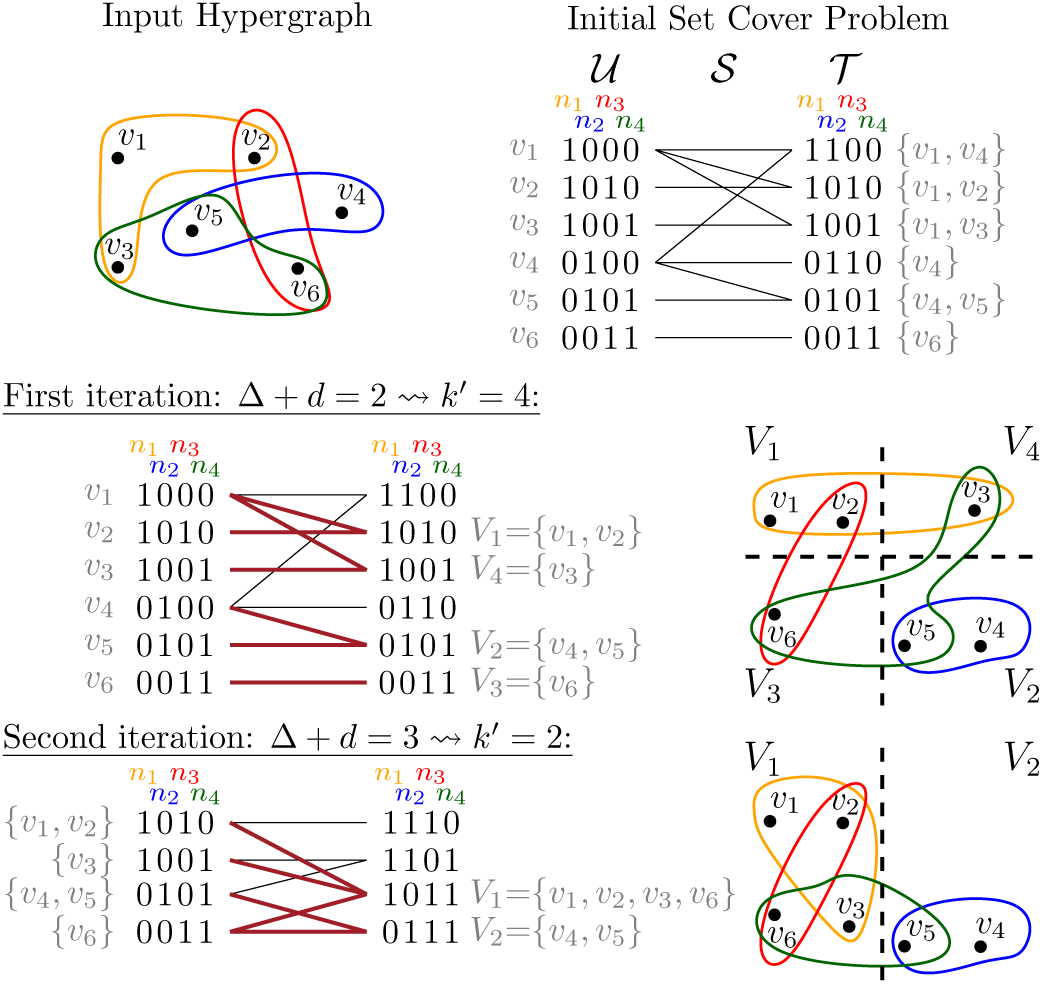
Modified example from Tan *et al.* [3]: A hypergraph with 6 vertices and 4 nets is partitioned into *k* = 2 blocks using their algorithm. The final partition has a cost of Δ + *d* = 3. Note that sets 𝒮_*j*_ are represented as *edges* in the bipartite graph.

#### Algorithm 1: Judicious Partitioning Algorithm of Tan et al. [3]

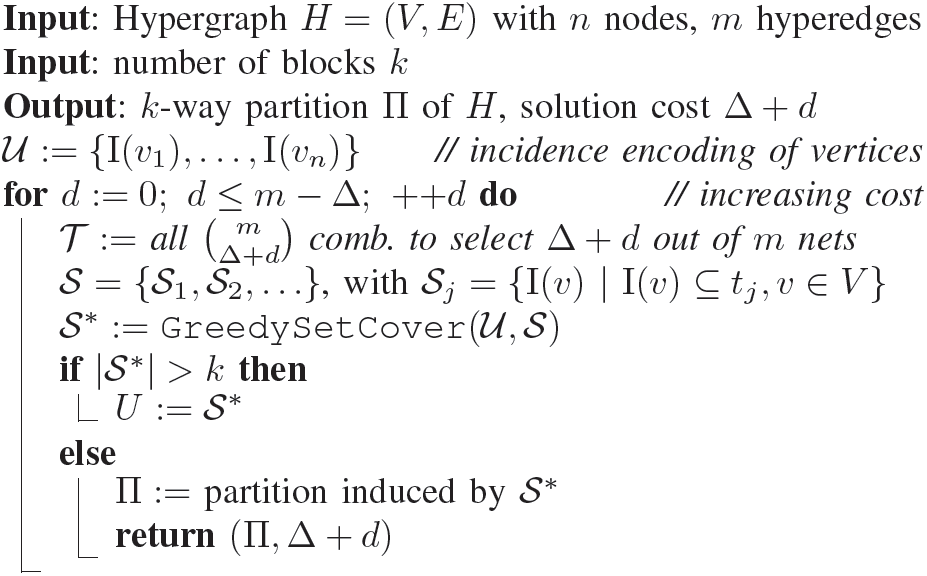

### D. Towards an Efficient Implementation

The algorithm might already require 𝒪 (*m*^Δ^) time for the first round, as 𝒮 can contain *all 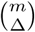* combinations. This is computationally infeasible, even for small hypergraphs.

#### a) Exploiting Instance-specific Features

However, we can exploit the fact that, by construction of our specific instance, *all* vertices have the same degree, that is, ∀*ν* ∈ *V* : *d*(*ν*) = Δ. Thus, all elements in 𝒰 are represented by exactly the same number of hyperedges. Therefore, it is possible to omit the first round (*d* = 0) of the algorithm as it will only group vertices *u* and *ν* that have the exact same incidence structure (i.e., I(*u*) = I(*ν*)). Instead, we perform one pass over the elements of 𝒰 and remove all duplicate entries (i.e., vertices with the same incidence structure).

In all subsequent iterations, the representations of elements in 𝒰 and 𝒮will only differ in *one* additional hyperedge. Thus, it is possible to generate all elements in 𝒮 by combining all pairs of elements in 𝒰. Hence, generating the initial collection of sets 𝒮 only requires 𝒪 (Δ*n*^2^) time, as we can compute the union of all pairs of elements in 𝒰 and keep exactly those where |𝒮_*j*_| = |𝒰_*j*_|+ 1. Since the greedy set cover algorithm always chooses the next set 𝒮_*i*_ such that it contains the largest number of *uncovered* elements, it is not necessary to compute all elements 𝒮_*i*_ with |𝒮_*i*_|= 1 a priori. Instead, these elements can be generated on-the-fly when the greedy algorithm has no elements with |𝒮_*i*_|≥2 left to process *and* has not found a solution that covers 𝒰. This approach substantially reduces the overall memory consumption, since generating 𝒮 naïvely would lead to 𝒪 (*nm*) 𝒮 _*i*_ elements with | 𝒮_*i*_|= 1, while our approach generates at most *n* such elements.

#### b) Improving Convergence Behavior

Henceforth, we denote such elements in 𝒮 that are generated on-the-fly and only cover one element *u* ∈𝒰 as *filler* elements. When the greedy algorithm has no elements |𝒮_*i*_|≥2 left and therefore adds a filler element to the solution 𝒮*, the block of the hypergraph partition corresponding to this filler element does not change. It will contain the same vertices as in the previous iteration, since filler elements only cover a single element of 𝒰. While this is not a problem in theory, it can substantially affect the convergence speed of the algorithm and hence overall running time. This is because with a large fraction of filler elements, the current number of blocks *k*′ converges only slowly to *k*, since many blocks will remain stable during subsequent iterations. Furthermore, this behavior can deteriorate solution quality, since more iterations will be required for merging two blocks and the solution cost (i.e., Δ+ *d*) increases with each iteration. By employing the greedy set cover algorithm, the judicious partitioning algorithm of Tan *et al.* [3] chooses an *arbitrary* filler element out of all combinations of Δ + *d* nets that constitute a superset of *u*. Since the chosen filler set 𝒮_*j*_ will be included in 𝒮* and thus becomes a new element of in the next iteration, some choices of 𝒮_*j*_ may be more difficult to cover using Δ + *d* + 1 nets than others. In the worst case, the element will again only be covered by another filler element with Δ + *d* + 1 nets.

To alleviate this issue, we deploy the following strategy when choosing a filler element for a yet uncovered element *u* ∈𝒰: We first compute the symmetric set difference between *u* and all other elements *u*′ ∈𝒰. We then choose the filler element 𝒮_*j*_ such that the difference to the closest element *u*′ decreases. This increases the chance that both, *u*, and *u*′ can be covered using a non-filler element in subsequent iterations. We will provide additional details on this in the paragraph after next.

#### c) Representing 𝒰 and 𝒮 Sets

Both, the elements of 𝒰, and the sets of 𝒮 comprise elements that are represented by hyperedges of the hypergraph. While Tan *et al.* [3] do not explicitly discuss how set elements are represented, their example (an enhanced version of which is shown in Figure 3) already hints at a potential implementation: each element can be represented as a *m*-bit bitvector, where a set bit at the *i*th position denotes that hyperedge *i* is part of the set element. Thus, set unions and intersections become bit-wise *and* and *or* operations, respectively. However, if *m* is large and Δ is small, these bitvectors become large and sparse and waste space as well as time. We therefore use a dedicated sparse bitvector representation where each element is explicitly represented by a set of hyperedge IDs. Evidently, there is a time and memory trade-off between the sparse and the bit-wise set operations and representations.

The two alternative representations need to support the following operations: (i) Compute the symmetric set difference between two sets, (ii) given two sets *a* and *b*, add an element from the relative complement *a*\*b* to *b* (supporting our strategy for handling filler elements as discussed above), and (iii) merge two sets in order to create new 𝒮 elements from two 𝒰 elements. In the dense bitvector representation (i) can be implemented by computing a `XOR`on the two bitvectors. The size of the set difference then corresponds to the number of set bits in the result (which can be efficiently counted using `popcount` instructions), (ii) amounts to simple bitwise operations, and (iii) corresponds to a bit-wise `OR`. In the sparse representation, (i) is implemented using a merge-based symmetric set difference, (ii) simply scans both sets and adds the corresponding element to the set, while (iii) amounts to a merge-based set-union operation. Note that for the sparse representations these operations require the sets to be sorted.

While the running times of all operations are constant for the dense representation, running times of the sparse representation slowly increase with growing set sizes in successive iterations of the algorithm. Given that we typically have to analyze large and comparatively sparse input instances, this performance degradation is offset by the acceleration of the initial iterations that are the most time consuming. In our experiments, the sparse representation performed better than the dense representation if set elements contained *<* 0.1% of the total elements. Also, the sparse representation outperformed the dense representation overall.

#### d) Putting Things together and Parallelization

In Algorithm 2 we provide the pseudo code for HyperPhylo. We generate the set 𝒮 by iterating over all pairwise combinations of elements in 𝒰, checking whether the union of the two elements creates a valid 𝒮 element (i.e.,|*u*_1_∪*u*_2_|= |*u*_1_|+ 1), and adding the new element to if the condition is fulfilled. Furthermore, we calculate the minimal distances between elements of 𝒰 and store a potential filler element for each element *u* ∈𝒰 in a map *d*_min_. Since there are no data dependencies between these operations, they can be performed in parallel for all pairs (*u*_1_, *u*_2_) ∈𝒰.

We then compute 𝒮* by first executing the greedy set cover algorithm sequentially until no more elements of 𝒮 can be used to cover 𝒰. In this case, we have to resort to filler elements in order to cover the remaining 𝒰 elements. Filler elements are created by adding an element *x*_*i*_ from the relative complement of *d_min*[*u*] and *u* to the element *u*. Again, this can be done in parallel for all yet uncovered elements of 𝒰.

##### Algorithm 2: HyperPhylo Judicious Partitioning

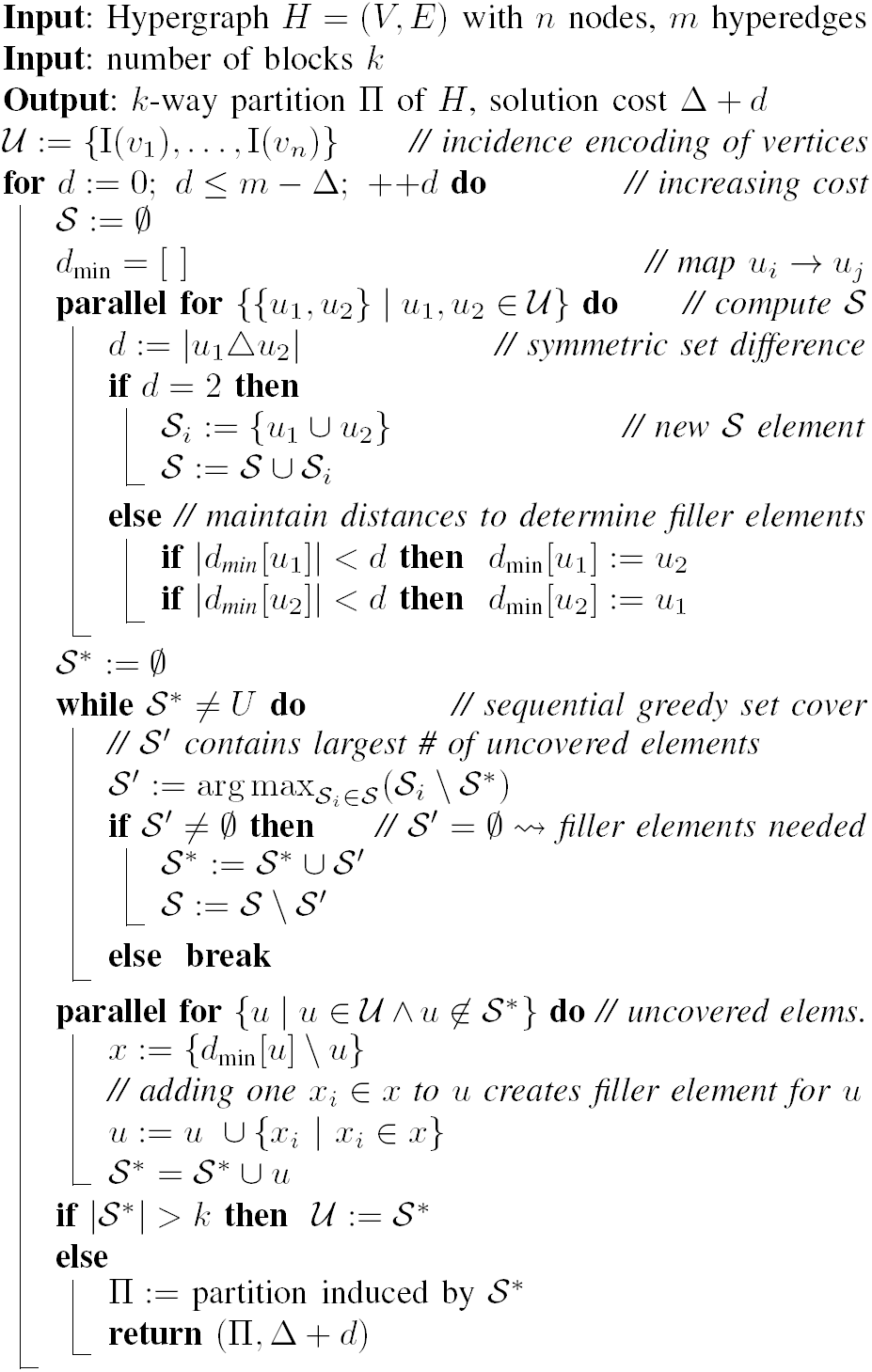

#### e) Implementation Details

Our dense set representation exploits x86 vector instructions through appropriate bit-vector starting address alignments in the memory allocation. Therefore, all loops that iterate over consecutive 64-bit blocks are vectorized automatically by the compiler.

In the greedy set cover algorithm, we need to repeatedly check if the entire set 𝒰 is already covered by 𝒮*. Instead of assessing if both sets, that is, the already covered elements of 𝒰 and 𝒰, are identical, it suffices to compare the corresponding set sizes.

The *parallel for* sections in Algorithm 2 are parallelized using OpenMP. As already mentioned, both parallel sections exhibit multiple possible execution paths resulting in a data-dependent running time performance. Therefore, we use the OpenMP dynamic workload scheduler to attain improved load balance.

The shared-memory data structures we use from the Intel® Threading Building Blocks library [14] are listed in Table I.

**TABLE I.**
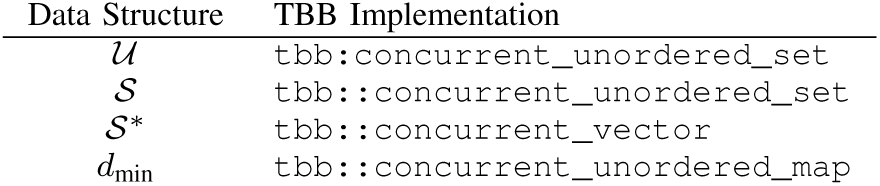
Intel® TBB Data structures used in our implementation.

## IV. EXPERIMENTAL RESULTS

HyperPhylo is written in `C++`and was compiled using `CLANG v.6.0.0` with the `-O3` flag. We also used OpenMP 3.1 [15] as well as the Intel^®^ Threading Building Blocks library version 2017.0 [14] for shared-memory parallelism.

### A. Hardware

We performed computational experiments on 3 distinct hardware platforms. Platform **A** has four 8-core Intel^®^ Xeon^®^ E5-4640 (Sandy Bridge) processors running at 2.4 GHz and 512 GB main memory. Platform **B** consists of two 16-core Intel^®^ Xeon^®^ E5-2683v4 (Broadwell) processors (2.1 GHz) and has 512 GB main memory. Platform **C** contains a 32-core AMD EPYC™ 7551P processor (2.0 GHz) and has 256 GB main memory. All platforms run Ubuntu 18.04.1 LTS.

### B. Verification

To increase our confidence that the implementation is correct, we verified its results on a small test case for which we explicitly computed the expected result using pen and paper. We also integrated an, as large as possible as well as reasonable, number of assertions into the code. Finally, we verified that all results represented a valid sites-to-cores mapping, that is, that each site of the input MSA is indeed assigned to exactly one core.

### C. Instances (MSAs)

For our experiments we use a total of four MSAs containing empirical sequence data from previous collaborative studies with biologists. The three smaller MSAs [16] comprise 59, 128, and 404 sequences, respectively. We will henceforth refer to these as *D59, D128*, and *D404*. *D59* has 7 MSA partitions and 6 951 sites in total, *D128* has 34 MSA partitions and 29 198 sites, and *D404* has 11 MSA partitions and 13 158 sites. From each of these four data sets, we extract both the smallest (referred to using suffix *-s* in Table II) and the largest (suffix *-l*) MSA partition. Furthermore, we use one substantially larger MSA, taken from the one thousand insect transcriptome evolution project [17], which we refer to as *supermatrix*. From this data set, which contains 50 MSA partitions and 413 459 sites in total, we selected four MSA partitions comprising 11 756, 20 753, 31 854, and 170 859 sites, respectively, with the last MSA partition being the largest in the entire *supermatrix* data set.

**TABLE II.**
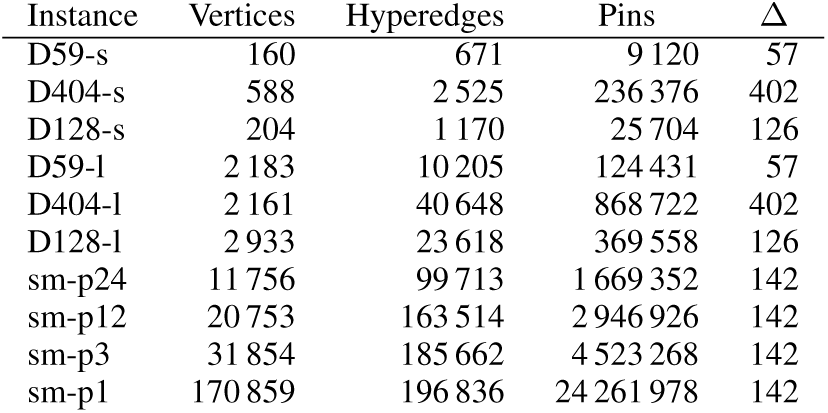
Properties of the hypergraphs generated from our data sets. D59-s represents the smallest MSA partition of data set *D59*, D59-l the largest etc. Sm-p1 is MSA partition 1 in the *supermatrix* data set. The number of sites of an instance is equal to the number of vertices of the corresponding hypergraph.

For each instance, we generated corresponding random trees and subsequently transformed them into hypergraphs as described in Section III-B. Note that each hypergraph corresponds to only *one* MSA partition of the respective MSA. The basic properties of the hypergraphs are provided in Table II.

### D. Parallel Scalability

To the best of our knowledge, there is no publicly available implementation of any judicious hypergraph partitioning algorithm. Therefore, we use the sequential implementation of HyperPhylo as a baseline for our speedup measurements. We demonstrate the scalability of our algorithm via the strong and weak scaling experiments summarized in Figures 4 and 5, respectively. Properties of the hypergraphs used in these experiments are summarized in Table III. We performed strong scaling experiments on all three hardware platforms. For strong scaling, the hypergraph is created by extracting a subset of 50 000 sites from the *supermatrix* data set. The corresponding hypergraph is then partitioned into *k* = 50 blocks (i.e., to distribute these sites onto *c* = 50 cores). The comparison of the dense and sparse representations in Figure 4 shows that the sparse representation performs better on all hardware platforms. We also observe that the sparse representation is more scalable than its dense counterpart as parallel efficiency of the dense representation drops noticeably when using more than one socket. Absolute running times for the strong scaling experiments are shown in Table IV. Note that these times are small compared to the times required for conducting production level inferences of phylogenetic trees under maximum likelihood on large clusters or supercomputers using hundreds or thousands of cores. Overall, when using all 32 cores of our three hardware platforms, the running time for judicious partitioning decreases by more than an order of magnitude. We generated data sets such that the number of sites divided by the number of cores used for computing the judicious partitioning remained constant at 5000 sites per core (e.g., 160 000 sites for 32 cores). The corresponding hypergraphs are always partitioned into *k* = 160 blocks. We use this setting because for a parallel PLF computation a core conducting likelihood calculations needs to be assigned at least 1 000 alignment sites (i.e., 160 000/1 000 = 160) for the PLF parallelization to scale. Note that Δ remains the same for all hypergraphs, as the same tree topology is used for all instances. The results depicted in Figure 5 show that our algorithm scales well on both platforms with an increasing number of threads. For small input sizes, the dense representation outperforms the sparse representation because it uses highly efficient vectorized bit operations, while the sparse representation employs element-wise integer comparisons. However, as the input size increases, the bit-wise encoding becomes less efficient. Therefore, the sparse representation performs better in these cases. For 160 000 sites, for example, each 𝒰 element and every element of a set 𝒮_*j*_ ∈𝒮 is encoded using 229 558 bits (all of which are potentially affected by the corresponding set operations), whereas the sparse representation only uses set operations on Δ+1 integers (i.e., the corresponding hyperedge IDs) in the first iteration.

**TABLE III.** Properties of the hypergraphs (derived from the *supermatrix* data set) used in weak and strong scaling experiments.

| # Sites | Scaling | Vertices | Hyperedges | Pins | $\Delta$ |
| --- | --- | --- | --- | --- | --- |
| 5 000 | weak | 5 000 | 57 631 | 710 000 | 142 |
| 10 000 | weak | 10 000 | 100 256 | 1 420 000 | 142 |
| 20 000 | weak | 20 000 | 132 093 | 2 840 000 | 142 |
| 40 000 | weak | 40 000 | 153 489 | 5 680 000 | 142 |
| 45 000 | weak | 45 000 | 158 230 | 6 390 000 | 142 |
| 50 000 | weak/strong | 50 000 | 163 381 | 7 100 000 | 142 |
| 60 000 | weak | 60 000 | 173 496 | 8 520 000 | 142 |
| 80 000 | weak | 80 000 | 192 543 | 11 360 000 | 142 |
| 85 000 | weak | 85 000 | 197 064 | 12 070 000 | 142 |
| 90 000 | weak | 90 000 | 200 072 | 12 780 000 | 142 |
| 100 000 | weak | 100 000 | 208 215 | 14 200 000 | 142 |
| 120 000 | weak | 120 000 | 200 967 | 17 040 000 | 142 |
| 125 000 | weak | 125 000 | 204 300 | 17 750 000 | 142 |
| 130 000 | weak | 130 000 | 207 048 | 18 460 000 | 142 |
| 140 000 | weak | 140 000 | 213 763 | 19 880 000 | 142 |
| 160 000 | weak | 160 000 | 229 558 | 22 720 000 | 142 |

**TABLE IV.** Absolute running times in minutes for strong scaling experiments on platforms **A, B**, and **C**.

| Threads<br># | Dense Representation |  |  | Sparse Representation |  |  |
| --- | --- | --- | --- | --- | --- | --- |
|  | A | B | C | A | B | C |
| 1 | 1368 | 789 | 629 | 746 | 1112 | 825 |
| 2 | 704 | 434 | 321 | 379 | 560 | 418 |
| 4 | 359 | 229 | 199 | 199 | 284 | 246 |
| 8 | 198 | 131 | 108 | 111 | 146 | 127 |
| 9 | 281 | 118 | 122 | 97 | 130 | 114 |
| 10 | 267 | 107 | 112 | 88 | 118 | 103 |
| 12 | 223 | 90 | 91 | 74 | 98 | 87 |
| 16 | 174 | 70 | 76 | 58 | 76 | 68 |
| 17 | 168 | 75 | 71 | 54 | 72 | 63 |
| 18 | 161 | 71 | 67 | 52 | 68 | 60 |
| 20 | 148 | 67 | 61 | 48 | 62 | 54 |
| 24 | 126 | 59 | 55 | 42 | 54 | 46 |
| 25 | 125 | 58 | 51 | 39 | 51 | 45 |
| 26 | 122 | 56 | 50 | 38 | 50 | 42 |
| 28 | 115 | 54 | 47 | 37 | 46 | 40 |
| 32 | 102 | 50 | 44 | 33 | 42 | 36 |

**Fig. 4.**
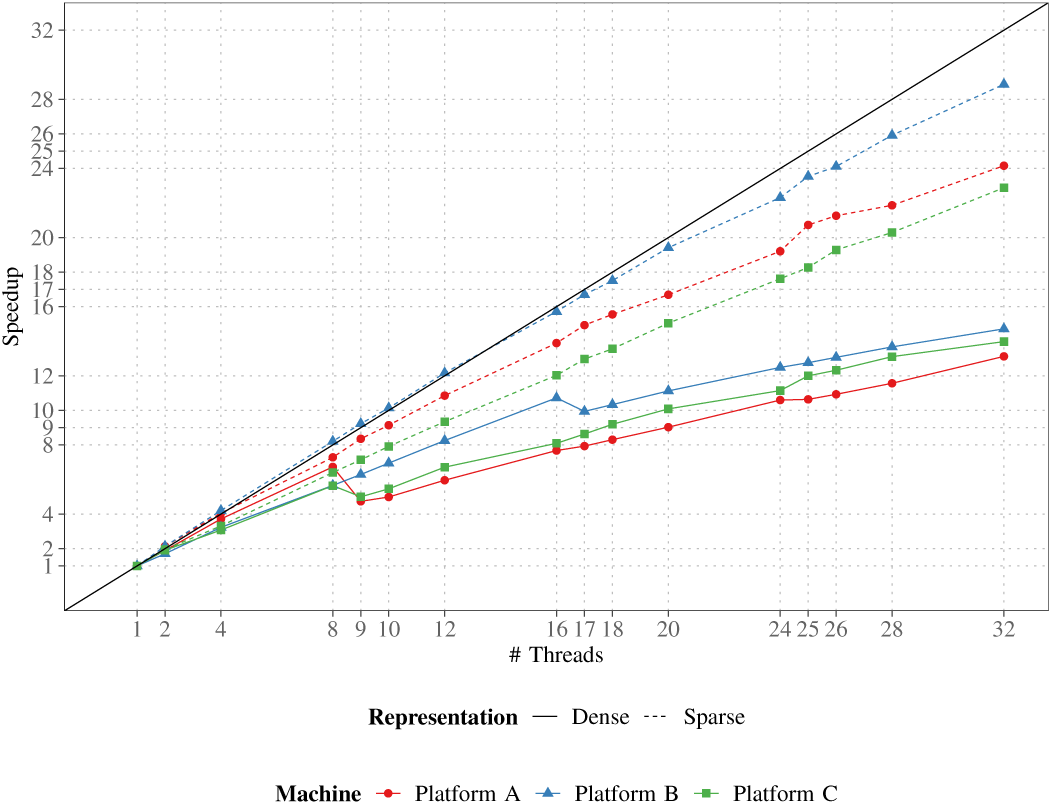
Speedup of HyperPhylo with increasing number of cores. The MSA instance contains 50 000 sites and the corresponding hypergraph is always partitioned into *k* = 50 blocks.

**Fig. 5.**
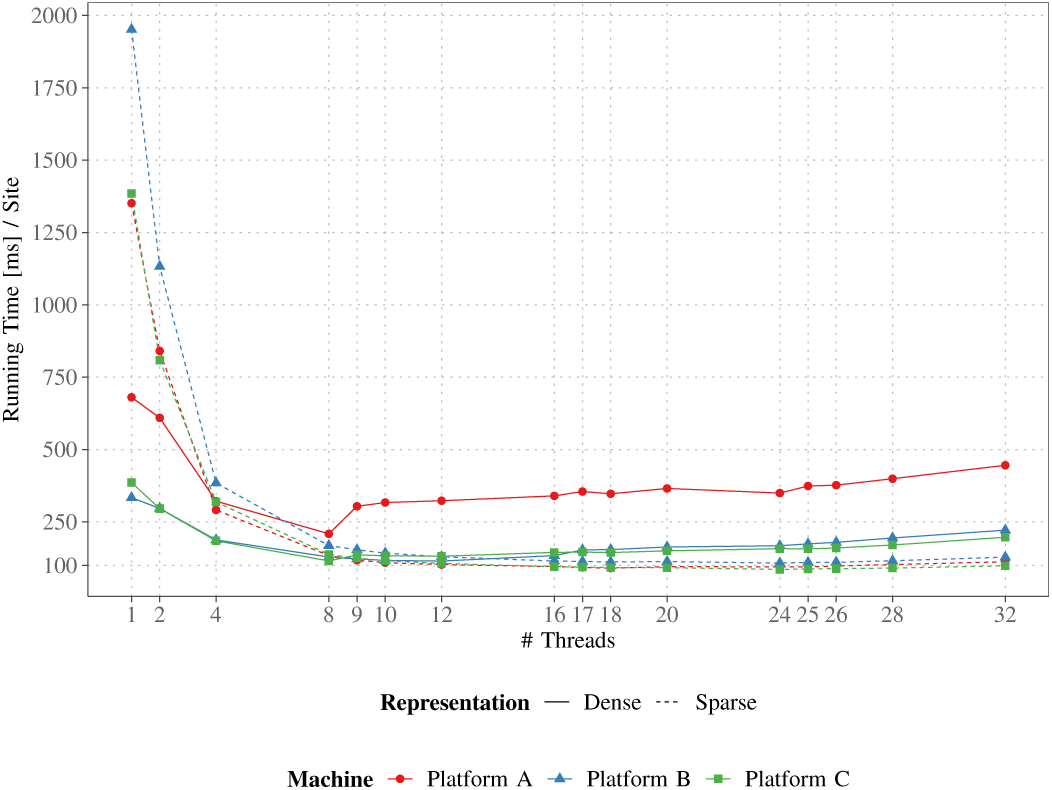
Weak scaling experiment with 5 000 sites per thread. The corresponding hypergraphs are always partitioned into *k* = 160 blocks.

### E. Solution Quality

To evaluate the quality of the phylogenetic data distribution of our HyperPhylo implementation compared to RDDA (the previous ad hoc algorithm), we computed data distributions for the MSAs described in Section IV-C on random trees generated with RAxML using both approaches. As solution quality measure, we use the ratio between *f* (*Z*) = max_*ζ∈Z*_ *flops*(*ζ*) and the lower bound. The lower bound is the value of *flops*() for a sequential calculation of the PLF, which is minimal as no repeats are lost, divided by the number of cores *c*. Thus, a quality measure value of 1 indicates that the solution is as good as the lower bound (i.e., optimal); a value of 2 means that the solution is two times worse than the lower bound etc. Figure 6 shows the average solution quality over all MSA partitions and over the number of cores *c* for RDDA and HyperPhylo. Since our quality measure is normalized by the lower bound, we use the geometric mean to appropriately average the ratios over all instances [18]. Detailed results for the distinct individual, single MSA partitions are shown in the Appendix.

**Fig. 6.**
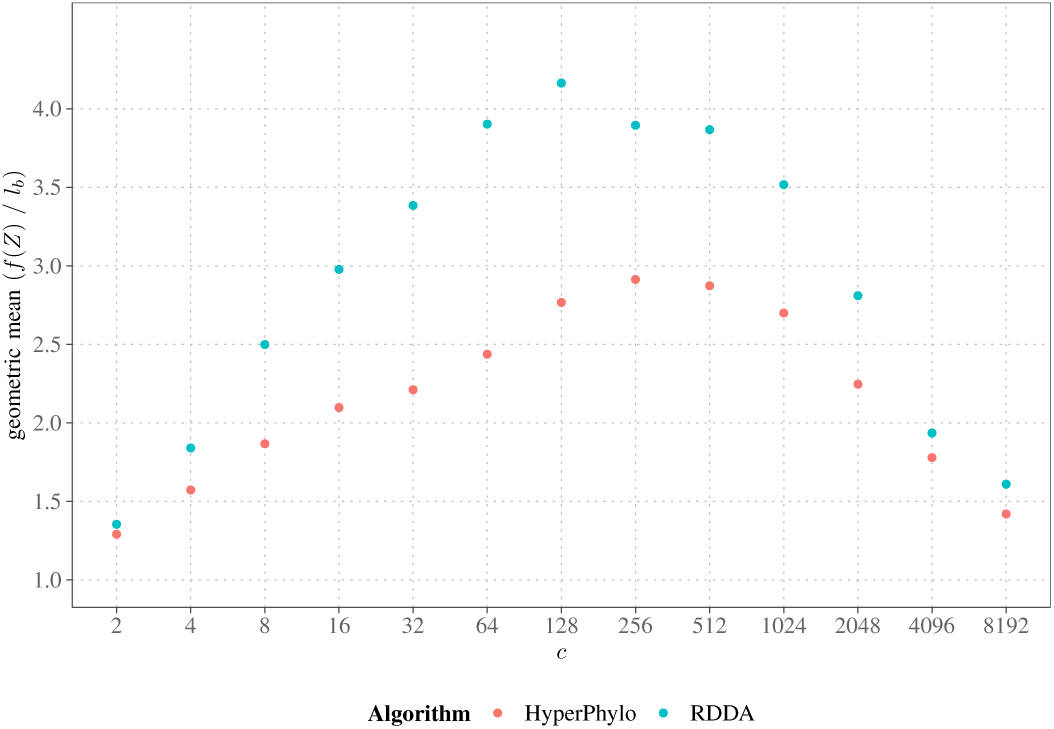
Average solution quality *f* (*Z*) = max_*ζ*∈*Z*_ *flops*(*ζ*) of the RDDA and HyperPhylo algorithms relative to the lower bound *l*_*b*_.

We observe that on average, HyperPhylo outperforms RDDA for all core counts. This is because RDDA does not take SRs into account when assigning the sites of a single MSA partition to two or more cores. In other words, the RDDA data distribution is more coarse grained as it only accounts for SRs at the MSA partition level.

In Figure 7, we show that the repeat loss (i.e., the number of SRs that are lost by assigning sites containing those SRs to distinct cores and which therefore lead to redundant computations on multiple cores) can be substantially reduced for data distributions on *c* = 2 up 8 192 cores (*k*-way partitions of the hypergraph). This shows that HyperPhylo computes data distributions which are better than those of RDDA in terms of *both* load balance *and* repeat loss.

**Fig. 7.**
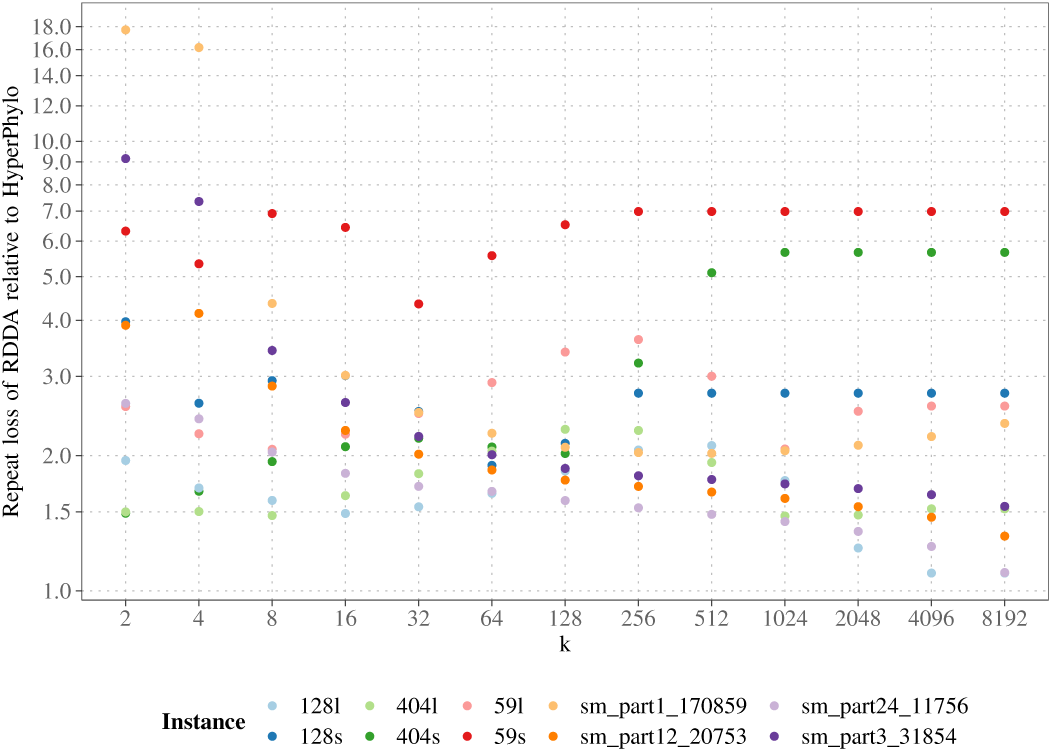
Repeat loss of RDDA relative to HyperPhylo. Note the log scale on the y-axis.

**Fig. 8.**
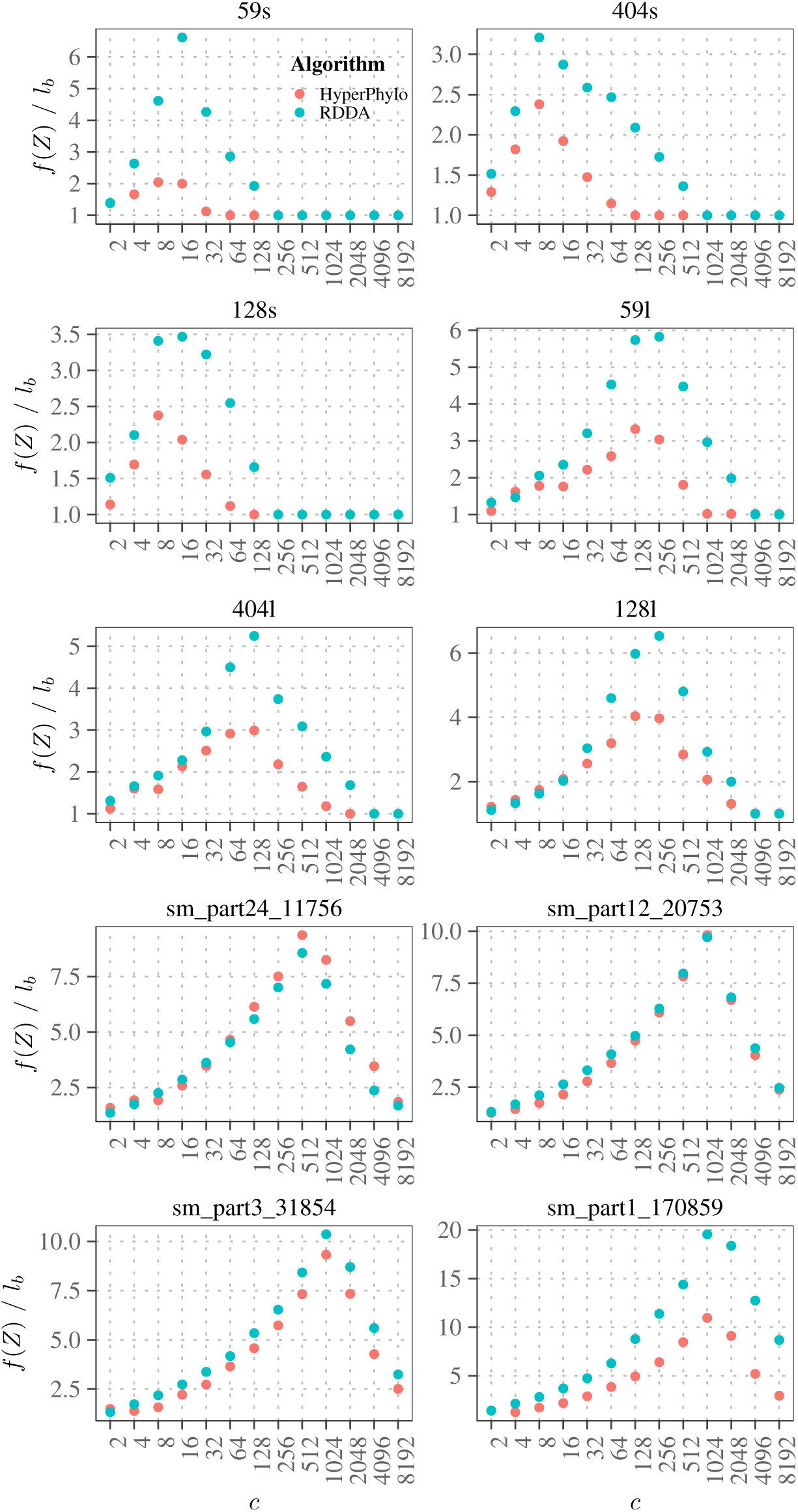
Per-instance solution quality *f* (*Z*) = max_*ζ*∈*Z*_ *flops*(*ζ*) of the RDDA and HyperPhylo algorithms relative to the lower bound *l*_*b*_.

## V. CONCLUSION

We have developed HyperPhylo, a highly efficient and scalable open-source implementation for a specific flavor of judicious hypergraph partitioning where every vertex has the same degree. The focus on this specific flavor is motivated by a real-world phylogenetics problem: optimal data distribution for massively parallel likelihood calculations with site repeats. We describe an algorithm and its efficient implementation for this specific version of judicious partitioning which we also make available as open-source code. Then, via weak and strong scaling experiments, we show that our algorithm exhibits good parallel efficiency. We also demonstrate that HyperPhylo can improve data distribution quality by up to 50% compared to the naïve algorithm. In addition, the repeat loss can be reduced by up to a factor of 18 compared to the naïve RDDA algorithm that can not take repeat loss into account.

While we suspect that the optimal phylogenetic data distribution problem under site repeats is NP-hard, we have thus far, not been able to devise a proof, mainly because of the unclear restrictions that the structure of the phylogenetic tree induces on any mapping. Further investigating this represents an avenue of future work.

In addition, we plan on developing an enhanced version of HyperPhylo that can dynamically switch between sparse and dense set representations depending on the current iteration and data set at hand.

In terms of practical deployment for phylogenetics, we consider that a hybrid implementation, that is, initially using the RDDA algorithm for assigning MSA partitions to cores and subsequently using HyperPhylo to assign sites to cores only for those MSA partitions that need to be split up among several cores, might work best. We intend to integrate this into our RAxML tool for maxmimum likelihood based phylogenetic inference.

## ACKNOWLEDGMENT

The authors wish to thank Matthias Wolf, Marcel Rademacher, and Sabine Kugler for discussions on the potential NP-hardness proof of the problem.

### APPENDIX

### DETAILED EXPERIMENTAL RESULTS

Note that for some instances, the number of cores becomes larger than the number of sites. In this case the assignment becomes trivial, which is why the quality measure is 1.

## Footnotes

1 available at https://github.com/lukashuebner/HyperPhylo

